# linearPOA: A parallel, memory-efficient framework for Partial Order Alignment with linear space complexity

**DOI:** 10.64898/2026.04.27.720899

**Authors:** Yanming Wei, Zhaoyang Huang, Pinglu Zhang, Qinzhong Tian, Yan Li, Quan Zou, Liang Yu

## Abstract

Multiple sequence alignment (MSA) is a fundamental problem in computational bioinformatics, playing a critical role in genome biology, especially in long read sequencing and assembly. One solution for representing and solving MSA is Partial Order Alignment (POA), which employs Directed Acyclic Graphs (DAGs) to represent sequence relationships. However, when facing the ultra-long, error-prone reads (e.g., >100 kbps), existing POA algorithms with quadratic space complexity become impractical due to excessive memory consumption. This paper introduces the linearPOA, which based on divide-and-conquer strategy to solve the POA, aimed at saving memory compared to quadratic space complexity algorithms like SPOA, abPOA and TSTA. Particularly notable is its capability to save up to 102.74 times memory usage when aligning sequences with 100 kbp reads, compared to the abPOA method using non-heuristic methods. The algorithm was implemented within the linearPOA library, providing functionality for POA and foundational support for sequencing analysis, like error correction for reads. The linearPOA algorithm provides memory-efficient algorithms for long-read sequencing, especially in directly assembling long reads like 100 kbp reads.

**Availability:** The linearPOA library is freely available at https://github.com/malabz/linearPOA, and the data underlying this article are available in Zenodo, at https://doi.org/10.5281/zenodo.15637837.

**Supplementary information:** Supplementary information are available at *BioRxiv* online.

## 1 Introduction

Multiple sequence alignment (MSA) is a fundamental problem in computational bioinformatics (Li and Liu, 2023; Li, et al., 2021; Lin, 2024; Liu, et al., 2023; Meng, et al., 2023; Wang, et al., 2023; Zhang, et al., 2022), playing a critical role in tasks such as including single nucleotide variant (SNV) calling (Li, 2012), protein function prediction (Wang, et al., 2024) and virus phylogenetic analysis (Chen, et al., 2023; Tang, et al., 2022; Wei, et al., 2020; Wei, et al., 2022). One solution for representing and solving MSA is Partial Order Alignment (POA) (Lee, et al., 2002), which employs Directed Acyclic Graphs (DAGs) to represent sequence relationships and applies Dynamic Programming (DP) to align sequences to the DAG - commonly referred to as sequence-to-graph alignment. The consensus sequence(Lee, 2003) generated by POA is widely used for analyzing long reads from sequencing platforms such as Oxford Nanopore Technologies (ONT) (Bowden, et al., 2019; Brown and Clarke, 2016) and Pacific Biosciences (PacBio)(Rhoads and Au, 2015). They play a vital role in downstream tasks, such as error correction and genome assembly of error-prone long reads (Gao, et al., 2021; Vaser, et al., 2017), which are essential in applications like single-cell analysis (Dai, et al., 2022; Fan, et al., 2020; Guo, et al., 2024; Huang, et al., 2024; Lebrigand, et al., 2020; Wang, et al., 2023; Wang and Yuan, 2024; Xie, et al., 2025; Xu, et al., 2025; Zhao, et al., 2024; Zhu, et al., 2024).

As modern multi-core processors become more widespread, parallelization techniques have become increasingly important for POA, leading researchers to focus more on accelerating POA process. SPOA (Vaser, et al., 2017) was the first approach to apply the Single Instruction Multiple Data (SIMD) to the POA process, significantly improving computational efficiency. abPOA (Gao, et al., 2021) enhanced SPOA by performing K-band(Zou, et al., 2012) in generating POA, which improved performance and reduced computational complexity. TSTA (Zong, et al., 2024) further improved performance by incorporating the active F-loop structure from BSAlign (Shao and Ruan, 2024), supporting both SIMD and multi-threading. Inspired by the wavefront alignment (WFA) algorithm (Marco-Sola, et al., 2023; Marco-Sola, et al., 2021), POASTA (van Dijk, et al., 2025) significantly improves POA performance, particularly in high-similarity alignment tasks. The sequence-to-graph alignment process is critical component of POA, and researchers widely explored the parallelization strategies to improve its performance. Several notable strategies have developed. PaSGAL (Jain, et al., 2019) focuses on parallelizing the alignment of multiple query sequences to reference graph, achieving high throughput in batch-processing scenarios. Astarix (Ivanov, et al., 2022) employs a parallelization strategy based on the A* algorithm, enhancing the efficiency of sequence-to-graph alignment through informed path exploration. Similarly, PVGwfa (Peng, et al., 2025) implements a parallel version of the graph wavefront alignment algorithm, offering accelerated performance in sequence-to-graph alignment tasks under the edit distance calculation model.

Pairwise sequence alignment (PSA) serves as the computational basis for POA. The Hirschberg algorithm (Hirschberg, 1975) proposed a linear space complexity solution through divide-and-conquer strategy. Building upon the Hirschberg algorithm, the Miller-Myers algorithm (Myers and Miller, 1988) developed based on Hirschberg algorithm, was developed to address the case of affine gap penalty in PSA. WFA2 (Marco-Sola, et al., 2023) extended divide-and-conquer strategy into WFA algorithm (Marco-Sola, et al., 2021), reducing its space complexity to linear. Inspired by the Four Russians algorithm (Arlazarov, et al., 1970), FORAlign (Wei, et al., 2025) implemented an cache-optimized algorithm to accelerate the Miller-Myers algorithm, achieving superior performance in low-similarity PSA.

However, to the best of our knowledge, several key challenges remain unaddressed: (a) There has been limited exploration of linear space complexity in POA algorithms. When facing the ultra-long, error-prone reads (e.g., >100 kbps), existing algorithms with quadratic space complexity become impractical due to excessive memory consumption; (b) Although the divide-and-conquer strategy is widely adopted in PSA, it has not been thoroughly investigated in the context of POA.

To overcome these limitations, we developed linearPOA, a lightweight, high-performance POA library based on the Hirschberg algorithm, designed for affine gap penalties. By employing a linear-space divide-and-conquer approach and parallelizing computation using the fork-join model (Lea, 2000), linearPOA achieves substantial improvements in performance and memory usage. Our implementation is especially effective for ultra-long reads, such as MPoX-simulated 100 kbp reads, outperforming traditional POA methods in both speed and memory efficiency.

## 2 Methods

In the following sections, we explain the key principles behind the linearPOA library. We begin by introducing the divide-and-conquer strategy of linearPOA. Then, we describe how this approach is efficiently parallelized to take advantage of modern multi-core architectures.

### 2.1 The framework of linearPOA

linearPOA is designed to address the computational challenges of POA by leveraging the Hirschberg algorithm in combination with several optimization strategies. We begin by introducing the concept of divide-and-conquer strategy for sequence-to-graph alignment which is the core operation in POA. Since POA is constructed through a series of sequence-to-graph alignment steps, optimizing this process is critical. To further enhance performance, we parallelize the alignment by applying parallelism to the divide-and-conquer function. While this approach significantly reduces memory usage, though at the expense of a modest runtime overhead. Figure 1 presents an overview of the linearPOA framework, illustrating how these strategies are integrated to improve both space efficiency and overall scalability of the POA process.

**Figure 1.**
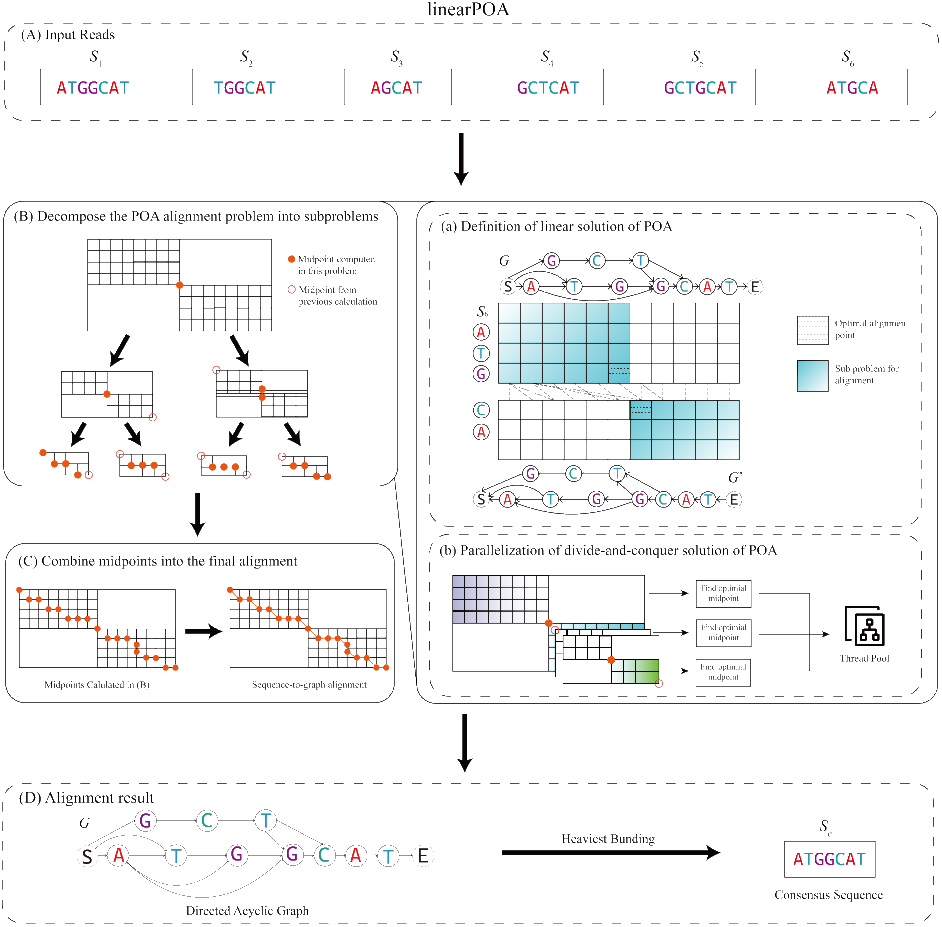
The flowchart of linearPOA. (A) The input is the unaligned sequences or reads. Firstly, the sequences are sorted, placing the longest sequence first. Each sequence is then aligned to the graph by sequence-to-graph alignment algorithm. The alignment process is divided into two key stages: (B) decompose the alignment into subproblems by identifying midpoints using divide-and-conquer strategy, and (C) sort the midpoints based on the topological order of the DAG and the order of the input sequence, concatenate the mid-points to construct the full alignment. The result of each alignment is used to incrementally insert the sequences into the graph. (D) Once all sequences have been processed, a complete DAG is constructed. A consensus sequence is then generated from this graph using the heaviest bundling algorithm (Lee, 2003). The decompose algorithm consists of two parts: (a) inspired by the Hirschberg algorithm, we define a linear space solution of POA. (b) To accelerate the process, we parallelize the divide-and-conquer algorithm by distributing the tasks into a thread pool. Each thread computes midpoints independently, allowing the overall process to be significantly accelerated

### 2.2 Divide-and-conquer strategy of linearPOA

In this section, we will briefly overview of the core concept behind linearPOA. The core base of linearPOA is parallelized Hirschberg algorithm (Hirschberg, 1975) applied on POA. We began with introducing the quadratic memory usage for affine gap penalty in solving POA. Next, we will delve into the core designation of linear memory usage algorithm.

#### 2.2.1 Quadratic memory solution for affine gap penalty POA

In this section, we introduce the traditional approach for solving POA with affine gap penalty scenario, which employed in many tools, like SPOA (Vaser, et al., 2017), abPOA (Gao, et al., 2021) and TSTA (Zong, et al., 2024). Let the sequence *S*_1_ = *S*_1,1_*S*_1,2_, … *S*_1,*M*_ has length *M*, and the DAG *G*_2_ has *N* nodes, the recurrence relations for affine gap penalty in POA is defined as shown in equations (1∼4):

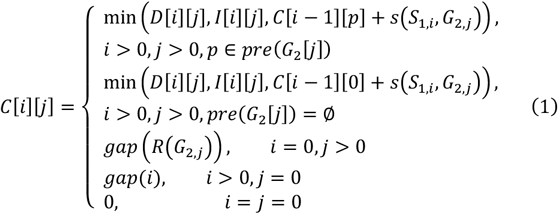

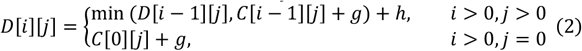

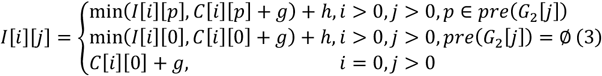

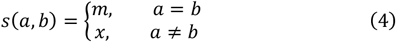

where *g* represents the gap open penalty, *h* denotes the gap extension penalty, and *ggp*(*x*) = *g* + *xh* defines the total gap cost for gap of length *x. R*(*N*) denotes the length of shortest path between node *N* in *G*_2_ and the virtual start node in *G*_2_. *C*[*i*][*j*] indicates the minimum alignment score between sequence *S*_1,1_, …, *S*_1,*i*_ and the subgraph of *G*_2_ which ending at node *G*_2,*j*_, considering all paths from start virtual node to *G*_2,*j*_, *D*[*i*][*j*] represents the minimum alignment score under the constraint that *S*_1,*i*_ is aligned to a gap, and *I*[*i*][*j*] signifies the case where *G*_2,*j*_ is aligned to a gap. It’s worth noting that, if *G*_2,*j*_ has no predecessor nodes, it’s obviously to be directly connected to virtual start node. node. In such cases, the corresponding values in the *I* and *C* matrices are initialized using column 0 of the dynamic programming matrix, which represents the alignment starting from the virtual start node. An example of sequence-to-graph alignment with quadratic memory is illustrated in Figure S1. The alignment score *s*(*a, b*) between character *a* and *b* is simplified by the match score *m* and mismatch score *x*. Obviously, solving the affine gap penalty via DP requires *O*(*MN*) time and space. To reduce space usage, we propose a linear space solution for POA. In the next section, we present the design of the linearPOA algorithm.

#### 2.2.2 Linear memory solution for affine gap penalty POA

To reduce space usage in POA, we designed an algorithm based on divide-and-conquer strategy, reducing the space complexity to linear. The score of our linear space algorithm lies in the method used to split the graph during alignment. Given a sequence *S*_1_ with length *M* and a DAG *G*_2_ with *N* nodes, obtaining the alignment with the minimum score under the affine gap penalty is achieved by identifying the midpoint (*i*^∗^, *j*^∗^) with the minimum score for *S*_1_ and *G*_2_. For consistency, the DAG has virtual start and end node, we denote them as *S* and *E*. The complete DAG *G*_2_ can be denoted as *G*_2,*S,E*_, representing the subgraph from the virtual start to the virtual end node.

Firstly, we define the reverse forms of the sequence *S*_1,*M*_ and the DAG *G*_2,*S,E*_. The reverse of *S*_1,*M*_ = *s*_1,1_, *s*_1,2_, …, *s*_1,*M*_ is denoted as 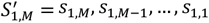, where *s*_1,*i*_ represents the *i*-th character in *S*_1_. For the DAG, we define *G*_2,*x,y*_ as the subgraph of *G*_2_ containing all paths from *g*_*x*_ to *g*_*y*_, where *g*_*x*_ is *x*-th node in *G*_2_. The reverse DAG 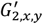 of the *G*_2,*x,y*_ is constructed by reversing all edges in *G*_2,*x,y*_, and the original start node *g*_*x*_ becomes the end node in 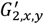 and the original end node *g*_*y*_ be the start node in 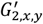. Next, we define the forward and backward phases for sequence-to-graph alignment. In the forward phase, we only need to consider the last row of the DP matrices *C* and *D*, resulting in vectors ***CC*** and ***DD***, based on the alignment of 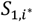 and *G*_2,*S,E*_. In the backward phase, we compute the last row of the reversed DP matrices *C*^*r*^, *D*^*r*^, denoted as ***CC***^*r*^ and ***DD***^*r*^, based on the alignment of 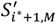 and 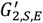.

Since ***DD***[0] and ***DD***^*r*^[0] are undefined, we simply set them as ***DD***[0] = ***CC***[0] and ***DD***^*r*^[0] = ***CC***^*r*^[0] for consistency. We now introduce the method for midpoint calculation. Based on the vectors defined above, consider the gap affine penalty model, we adopt the design strategy of the Hirschberg algorithm. Each time when searching for the minimum score coordinates in *S*_1,*M*_ and *G*_2,*S,E*_, fixing *i*^∗^ = ⌊*M*/2⌋, solve for *j*^∗^ using equation (5):

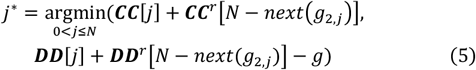

where *npxn*(*g*_2,*j*_) denotes the successor nodes of node *g*_2,*j*_ in the DAG. The key difference between the traditional Hirschberg algorithm and our proposed method lies in the structure of the alignment cases: Hirschberg aligns linear sequences with single-predecessor and single-successor characters. In contrast, our algorithm supports the DAG structure, enabling alignment of nodes with multiple predecessors and successors. After identifying the minimum score at the fixed row, our algorithm performs a case-by-case analysis:

1. Type 1 midpoint: If the minimum score comes from ***CC***[*j*] + ***CC***^*r*^*N* − *npxn*(*g*_2,*j*_), the problem is divided into solving (i) the minimum score for 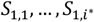 and all paths from the start node to 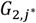 and (ii) the minimum score for 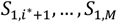 and the successor node 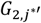 where 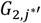 is the successor of 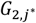 with the minimum alignment score;
2. Type 2 midpoint: If the minimum score comes from ***DD***[*j*] + ***DD***^*r*^*N* − *npxn*(*g*_2,*j*_) − *g*, the problem is divided into solving the minimum score for *S*_1,1_, …, 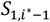 and all nodes from the start node to 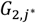, and (ii) the minimum score for 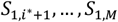 and the successor 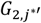 which the successor 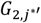 has the minimum score in all of the successors of 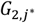.

Figure 2 illustrates the Type 1 and Type 2 midpoint scenarios in our algorithm. This recursive process continues until the sequence length or the number of DAG nodes is reduced to less than 1, at which point boundary cases are applied. The pseudo code for the linearPOA algo rithm is shown in Algorithm S1.

**Figure 2.**
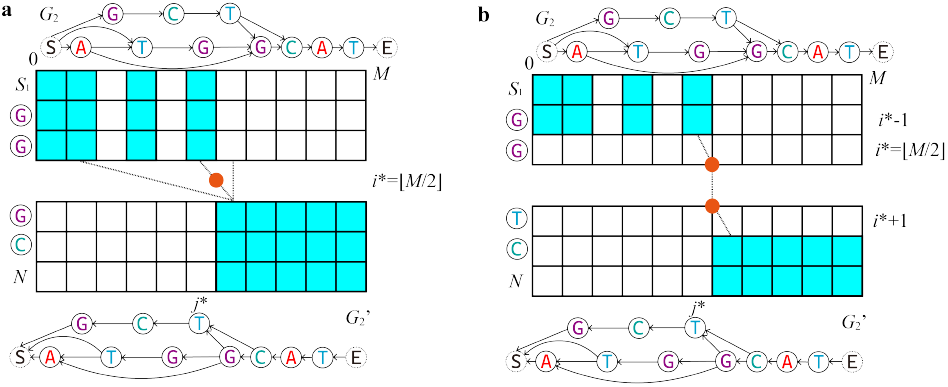
Divide-and-conquer strategy in linearPOA under the affine gap penalty model. To reduce space complexity, linearPOA fixes row index *i*^∗^ = ⌊*M*/2⌋, and finds the minimum score in two cases: (a) Simply consider the minimum score from the upper-left and lower-right parts (type 1 midpoint); (b) Consider the minimum score with upper-left and lower-right parts, but align 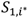 and 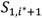 with gaps (type 2 midpoint). It’s worth noting that, the DAG are topologically sorted, allowing the subproblems to be cleanly divided according to topological order. The colored regions in the DP matrix represent the sub-problem blocks. When dividing the problems, the dependencies between subproblems must be carefully recalculated to ensure continuity in the alignment path across recursive steps

In summary, the linearPOA solved the POA problem by divide-and-conquer strategy. The algorithm divides the alignment task into subproblems based on two types of midpoints, which identify the optimal coordinates corresponding to the minimum alignment score. The next section will discuss the parallelization strategy used in linearPOA to further enhance performance.

## 3 Experiments and Results

### 3.1 Datasets and measurement

In this section, we conduct separate evaluations with linearPOA and compare its performance against the modern POA methods, like SPOA, abPOA and TSTA. For more and all-to-all comparison, we designed the simulation experiment and real experiment. This part will introduce the experimental design in our comparisons. For fully compare between our method and other POA methods, we designed the simulation experiment and real experiment. We briefly introduce the experimental design for our comparison.

#### 3.1.1 Simulated Datasets and methods

POA can be used in many cases, like the long-read error correction (Vaser, et al., 2017). To explore the advantages of space complexity of linearPOA and simulate the extremely long-read cases, we collected sequences from the SARS-CoV-2 dataset, which collected from GISAID (Shu and McCauley, 2017), the mitochondrial genomes (mt) (Tanaka, et al., 2004; Tang, et al., 2022; Wei, et al., 2022), the GMGC dataset (Coelho, et al., 2022), the HIV dataset (https://www.hiv.lanl.gov/), and the MPoX dataset (Ma, et al., 2022). We chose one sequence in every dataset as the reference sequence, the length of selected sequences are calculated by seqkit (Shen, et al., 2016), is shown in Table 1. Next, we generate different sequencing depths (3×, 5×, 10×, 30× and 50×) of these sequences by modified PBSIM2 (Ono, et al., 2021), which PBSIM2 only generate the forward strand sequences. We use the models provided in PBSIM2, named P6C4 and R103, to simulate the sequencing model for PacBio and ONT. Each depth and sequence repeatedly run 10 times to generate 10 files, and these files were aligned by the determined methods, like abPOA, TSTA and linearPOA, to generate a consensus sequence. We did not test against POASTA, PaSGAL, Astarix and PVGwfa since these methods are unable to generate consensus sequence. In our experiments, all method parameters are determined by affine gap penalty were set as *m* = 0, *x* = 4, *g* = 6, *h* = 2 (denoted as 0,4,6,2) and *m* = 0, *x* = 6, *g* = 5, *h* = 3 (denoted as 0,6,5,3). All tests are running in AMD EPYC 7H12 Processors, using 96 cores with 256 GB memory. The testing methods are shown in Table 2.

**Table 1.** The length of selected sequences in simulated dataset. Due to the limitation of PBSIM2, we cut the selected MPoX sequence to 100,000 bps.

| Reference | Dataset name | Length of selected sequence |
| --- | --- | --- |
| HIV | HIV | 10,372 |
| Mt | mt | 16,578 |
| SARS-CoV-2 | SARS-CoV-2 | 29,927 |
| GMGC | GMGC | 40,005 |
| MPoX | MPoX-100000 | 100,000 |

**Table 2.**
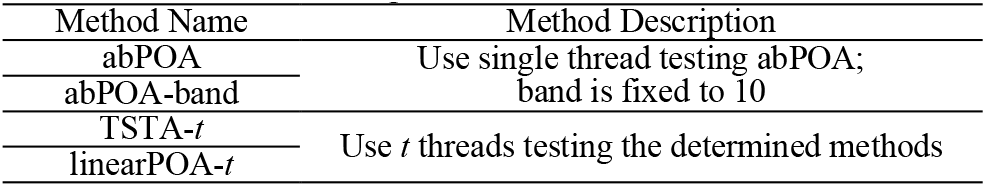
Testing methods in simulated datasets Method Name Method Description.

#### 3.1.2 Real Datasets and methods

To evaluate the performance of our method in the real-life scenarios, we modified the popular long-read error correction method Racon(Vaser, et al., 2017), to replace the consensus generation step. We ignore the window information provided by Racon, and directly align window sequences. The test is running in the error correction mode on real datasets listed in Table 3. We tested the following methods: SPOA, abPOA, abPOA-band, TSTA, and linearPOA, for consensus sequence generation within the Racon (Vaser, et al., 2017) pipeline. To ensure a fair comparison and eliminate potential bias introduced by Racon, the window information provided by Racon was disabled for all methods. For the corrected reads, we mapped them to the genome by minimap2 (Li, 2018). We then used miniasm (Li, 2016) for simply assembling correct reads. The alignment arguments and testing processors which used in real datasets are same as simulated datasets.

**Table 3.** Details of real datasets.

| Genome name | Genome size | Sequencing Technology | Number of sequences | Depth | Dataset Source |
| --- | --- | --- | --- | --- | --- |
| *Lambda phage* | 48,502 | ONT R7.3 | 646 | 113 | Racon |
| *E. Coli* K-12 | 4,641,652 | ONT R7.3 | 25,483 | 54 |  |
|  |  | PacBio HiFi | 6,305 | 20 | PacBio website |

#### 3.1.3 Experimental Metrics

To ensure a comprehensive assessment of multithreaded results, we monitored the utilization of CPU cores during testing. Our analysis focused on time taken for computation, space utilization and similarity between reference sequence and consensus sequence calculated with FORAlign (Wei, et al., 2025), WFA2 (Marco-Sola, et al., 2023) and Minimap2 (Li, 2018) in simulated datasets, mapped pairs to corrected sequences and mapped assembled corrected sequences to genome by miniasm (Li, 2016) in real datasets. We employed speedup metrics to measure the performance of different methods and the speedup, which are defined by equations (8) and (9):

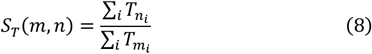

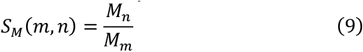

Where *m* and *n* represent different methods, 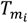 denotes the time usage for method *m* in test case *i*, and *M*_*m*_ indicates the maximum memory usage of method *m*. The similarity between reference sequence *p* and consensus sequence *c* are defined by equation (10):

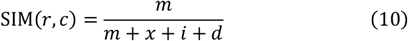

Where *m* represent match characters between *p* and *c, x, i, d* represent mismatch, insertion and deletion characters between *p* and *c*, respectively. Same as before, the similarity is the average of test cases.

### 3.2 Results

In this section, we present the experimental results that demonstrate the advantages of linearPOA. Our evaluation can be divided into the simulated datasets and the real datasets. In simulated datasets, we mainly demonstrate the advantages memory efficiency and runtime performance of linearPOA, as well as the effectiveness of its parallelization strategy. In real datasets, we showcase the practical applicability of linearPOA in the context of third-generation sequencing technologies.

#### 3.2.1 Experimental Results on simulated dataset

In this section, we compare linearPOA with abPOA and TSTA by evaluating the time and memory during the consensus sequence generation process on simulated data, and assess the quality of consensus sequence.

We then delve into our method, examining the effectiveness of its divide-and-conquer parallelization and the impact of different parameter settings on performance.

##### Time and Memory Comparison between linearPOA, abPOA and TSTA

We tested the time consuming and memory usage between our method and other methods, and the result is summarized in Table 4 and Table 5. As shown in Table 4, linearPOA demonstrates significantly lower memory consumption across all datasets, particularly in large-scale cases such as GMGC and MPoX-100000. In these scenarios, other methods fail to construct the DAG and generate consensus sequence due to excessive memory requirements. While linearPOA consumes more computation time to achieve this reduction in memory usage, the trade-off is especially beneficial for long-sequence datasets. Table 5 shows that the consensus sequence quality produced by linearPOA is comparable to or better than that of other methods, confirming that the space saving strategy does not compromise the accuracy of POA results.

**Table 4.**
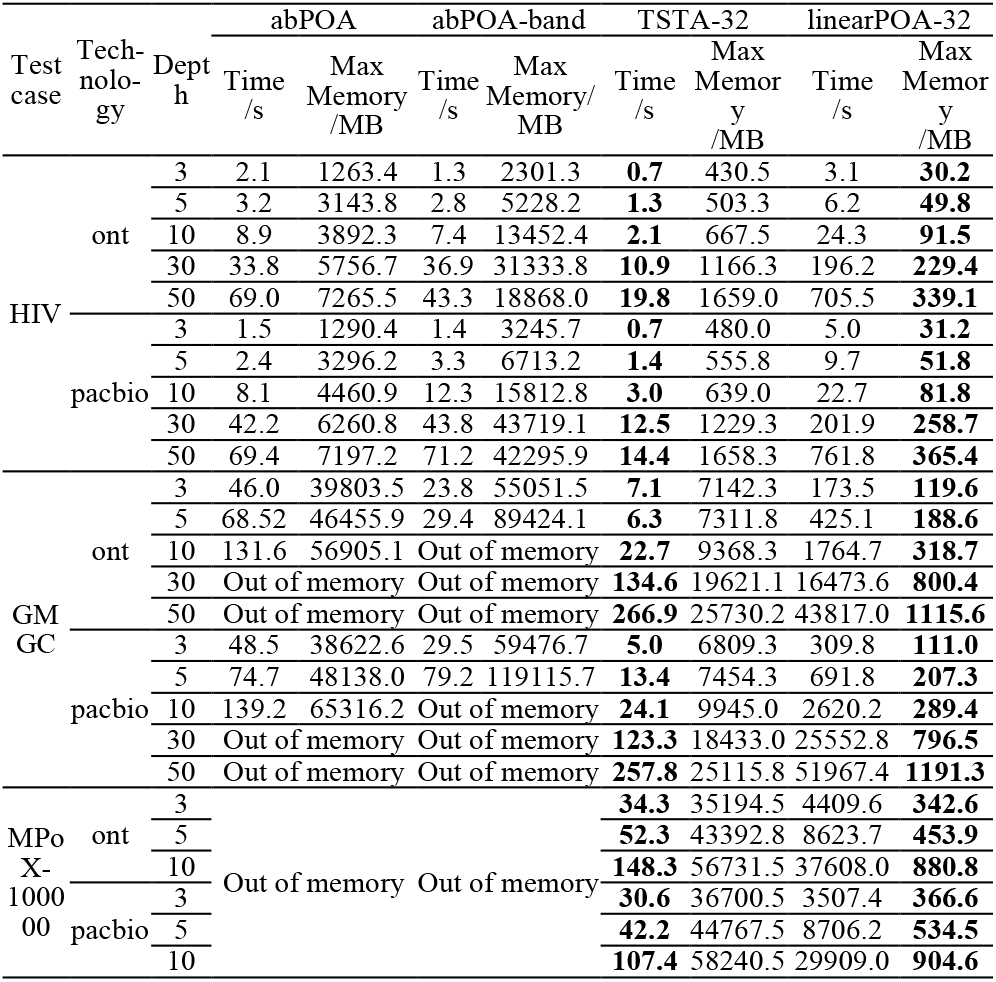
Time, memory and similarity between different methods in simulated dataset, tested in arguments 0,4,6,2. Due to only linearPOA-32 can run MPoX-100000 30× and 50× dataset, we do not show the results for these datasets in this table.

**Table 5.** Similarity results in simulated dataset, tested in arguments 0,4,6,2. “-” in this table means tested sequence has multiple mapped to reference.

| Test case | Technology | Depth | abPOA |  |  | abPOA-band |  |  | TSTA |  |  | linearPOA |  |  |
| --- | --- | --- | --- | --- | --- | --- | --- | --- | --- | --- | --- | --- | --- | --- |
|  |  |  | FORAlign similarity | WFA2 similarity | minimap2 similarity | FORAlign similarity | WFA2 similarity | minimap2 similarity | FORAlign similarity | WFA2 similarity | minimap2 similarity | FORAlign similarity | WFA2 similarity | minimap2 similarity |
| HIV | ont | 3 | 92.37% | 92.37% | 94.99% | 82.55% | 82.13% | - | 92.66% | 92.65% | 95.24% | 92.84% | 92.84% | 95.30% |
|  |  | 5 | 92.84% | 92.83% | 94.40% | 81.93% | 81.58% | - | 93.85% | 93.82% | 95.41% | 94.31% | 94.31% | 95.49% |
|  |  | 10 | 95.13% | 94.96% | 95.55% | 93.95% | 93.61% | - | 95.44% | 95.41% | 95.94% | 94.94% | 94.94% | 95.27% |
|  |  | 30 | 96.11% | 96.11% | 96.32% | 91.08% | 91.06% | 96.18% | 96.31% | 95.88% | 96.29% | 93.29% | 93.29% | 93.56% |
|  |  | 50 | 96.31% | 96.31% | 96.48% | 88.12% | 88.01% | 96.41% | 96.12% | 95.93% | 96.30% | 92.76% | 92.76% | 93.24% |
|  | pacbio | 3 | 86.44% | 86.43% | 91.24% | 80.19% | 79.68% | - | 87.53% | 87.50% | 92.35% | 87.91% | 87.90% | 92.47% |
|  |  | 5 | 88.19% | 88.18% | 91.19% | 73.93% | 73.83% | - | 90.01% | 89.96% | 93.11% | 90.11% | 90.08% | 92.72% |
|  |  | 10 | 90.96% | 90.94% | 92.54% | 90.10% | 90.10% | 85.52% | 92.13% | 92.04% | 93.65% | 91.47% | 91.45% | 92.56% |
|  |  | 30 | 92.53% | 92.54% | 93.51% | 92.55% | 92.22% | 93.74% | 92.86% | 92.76% | 93.88% | 90.96% | 90.88% | 91.75% |
|  |  | 50 | 92.71% | 92.70% | 93.62% | 88.22% | 87.98% | - | 92.79% | 92.46% | 93.83% | 87.08% | 87.08% | 88.25% |
| GMGC | ont | 3 | 93.40% | 93.39% | 96.64% | 84.79% | 82.35% | - | 93.72% | 93.67% | 96.51% | 92.38% | 92.37% | 95.15% |
|  |  | 5 | 94.45% | 94.38% | 97.81% | 90.00% | 89.90% | - | 95.22% | 95.21% | 98.06% | 94.29% | 94.29% | 95.30% |
|  |  | 10 | 94.97% | 94.97% | 98.93% |  | Out of memory |  | 95.63% | 95.60% | 98.72% | 94.79% | 94.79% | 95.11% |
|  |  | 30 |  | Out of memory |  |  | Out of memory |  | 96.06% | 95.90% | - | 84.82% | 84.81% | 84.99% |
|  |  | 50 |  | Out of memory |  |  | Out of memory |  | 96.30% | 96.25% | 99.21% | 93.61% | 93.61% | 93.99% |
|  | pacbio | 3 | 91.82% | 91.82% | 94.73% | 79.10% | 76.79% | - | 91.70% | 91.70% | 94.94% | 88.54% | 88.53% | 92.70% |
|  |  | 5 | 93.67% | 93.52% | 94.72% | 73.01% | 70.73% | - | 92.40% | 92.37% | 96.04% | 90.35% | 90.35% | 93.08% |
|  |  | 10 | 94.64% | 94.64% | 95.10% |  | Out of memory |  | 93.56% | 93.43% | 98.06% | 91.74% | 91.73% | 92.67% |
|  |  | 30 |  | Out of memory |  |  | Out of memory |  | 93.60% | 93.45% | 98.32% | 90.97% | 90.96% | 91.69% |
|  |  | 50 |  | Out of memory |  |  | Out of memory |  | 93.67% | 93.50% | 98.24% | 87.94% | 87.95% | 88.95% |
| MPoX-100000 | ont | 3 |  |  |  |  |  |  | 94.15% | 94.13% | 96.95% | 92.81% | 92.81% | 95.28% |
|  |  | 5 |  |  |  |  |  |  | 94.42% | 94.39% | 97.41% | 93.75% | 93.75% | 95.08% |
|  |  | 10 |  |  |  |  |  |  | 95.76% | 95.75% | 98.86% | 94.84% | 94.84% | 95.19% |
|  |  | 30 |  |  |  |  |  |  |  | Out of memory |  | 94.58% | 94.57% | 94.81% |
|  |  | 50 |  |  |  |  |  |  |  | Out of memory |  | 93.32% | 93.31% | 93.75% |
|  | pacbio | 3 |  | Out of memory |  |  | Out of memory |  | 88.11% | 88.06% | 92.80% | 88.17% | 88.16% | 92.70% |
|  |  | 5 |  |  |  |  |  |  | 89.95% | 89.75% | 93.06% | 89.31% | 89.31% | 91.89% |
|  |  | 10 |  |  |  |  |  |  | 91.69% | 91.64% | 93.36% | 90.59% | 90.60% | 91.81% |
|  |  | 30 |  |  |  |  |  |  |  | Out of memory |  | 90.42% | 90.35% | 91.20% |
|  |  | 50 |  |  |  |  |  |  |  | Out of memory |  | 88.95% | 88.96% | 90.11% |

##### Performance Evaluation of Divide-and-Conquer Parallelization in linearPOA

To thoroughly evaluate the speedup parallelization performance of linearPOA, we tested linearPOA using different numbers of threads and measured the corresponding execution time and memory usage. The results are shown in Figure S2 and Figure S3, covering all-simulated datasets. As shown in these figures, linearPOA demonstrates strong scalability, the execution time decreases as the number of threads increases. This indicates that the divide-and-conquer strategy used in linearPOA can be effectively parallelized across multiple cores, although the speedup tends to saturate earlier due to limited task granularity. It is also worth noting that while performance improves with more threads, memory usage also increases. This is expected because each thread maintains its own intermediate data structures during the alignment process. The trade-off between time and memory is a key consideration: linearPOA uses additional memory to achieve faster computation through parallelism. Nevertheless, the memory consumption remains manageable, especially when compared to other POA methods, and is significantly lower in multithreaded runs due to the algorithm’s linear-space design.

##### Performance Evaluation of alignment parameter settings in linearPOA

The alignment parameter settings are another important factor that influences the quality of the consensus sequence. To evaluate this effect, we tested all methods using two different sets of scoring parameters: 0,4,6,2 and 0,6,5,3, and the results are shown in Table S1. From the results, it is evident that the choice of parameters affects the outcome of the consensus generation process. Variations in scoring systems can lead to differences in both accuracy and error rates of the final consensus sequence, highlighting the importance of selecting alignment parameters based on the characteristics of the input data.

#### 3.2.2 Experimental Results on real dataset

In this section, we present a detailed analysis of the real dataset results, as shown in Figure 3. We observe that using a smaller window size generally improves the quality of the assembled genome across most cases. Our method linearPOA used significantly less memory compared to other approaches, only a slight reduction in consensus quality and a slower runtime. This trade-off is particularly valuable when larger window sizes are required during the sequence correction process, where traditional methods often fail to handle the memory demands of large-scale POA computations. It’s important to note that, our modified methods do not utilize the window information provided by Racon, which limit the achievable similarity in the consensus results. Consistent with the findings from the simulated dataset, the alignment parameter settings also have impact on the quality of the consensus sequence (see Table S2 in Supplementary Material for detailed results).

**Figure 3.**
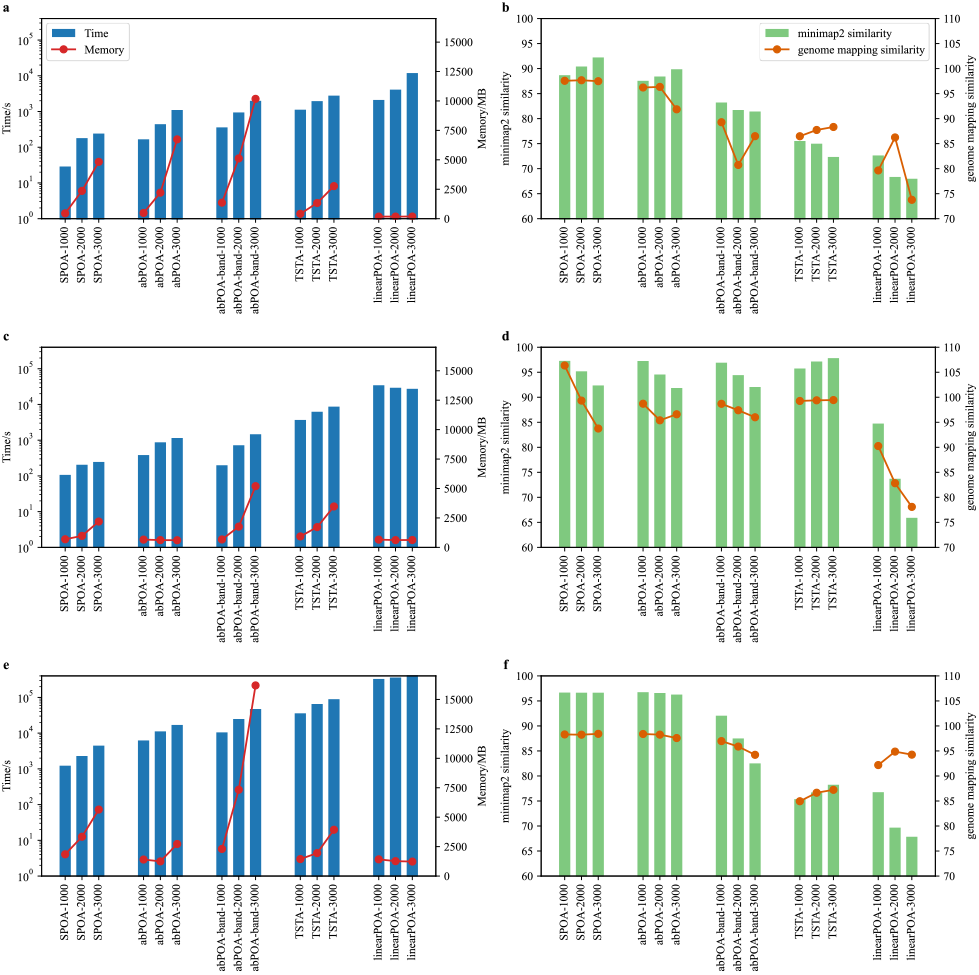
Performance results on real datasets using alignment parameters 0,4,6,2. (a) Runtime and memory usage for the *Lambda phage* dataset. (b) Similarity of corrected sequences for the *Lambda phage* dataset. (c) Runtime and memory usage for the *E. coli* K-12 dataset sequenced with HiFi technology. (d) Similarity of corrected sequences for the *E. coli* K-12 HiFi dataset. (e) Runtime and memory usage for the *E. coli* K-12 dataset sequenced with ONT technology. (f) Similarity of corrected sequences for the *E. coli* K-12 ONT dataset.

## 4 Conclusion

In this study, we introduced linearPOA, a parallelized library that implies the Hirschberg algorithm to support linear space complexity affine gap penalty alignment in POA. The efficiency of linearPOA is primarily influenced by the lengths of the sequences, consistently demonstrates stable speedup ratios, which makes it an effective solution for consensus sequence generation process. Compared to the traditional quadratic space complexity series methods, linearPOA demonstrates superior performance, particularly when aligning long reads like MPoX. The long reads were used in third-generation sequencing analysis, and our method plays a crucial role in supporting this analysis, where traditional methods often limited to deliver results without proper optimization. Our development efforts have culminated in linearPOA library compatible with both Linux and Windows platforms.

In future development, we aim to expand the capabilities of the linearPOA library to support 2-piece affine gap penalty parameters and various scoring matrix configurations. Additionally, we aim to adopt optimization strategies from TSTA to further accelerate the linearPOA process. Furthermore, we plan to add more functions in linearPOA, including heuristic methods, DAG format to FASTA format and related approaches. The linearPOA package is freely available at https://github.com/malabz/linearPOA. It has been tested on the Linux-like and Windows operating system. The data underlying this article are available in Zenodo, at https://doi.org/10.5281/zenodo.15637837.

## Acknowledgements

We acknowledge the help from the other group members: Yixiao Zhai, Tong Zhou, and Yizheng Wang for providing critical opinions during the preparation.

## Funding

This work has been supported by the National Natural Science Foundation of China (Nos: 62472344 and 62452107), the Xidian University Specially Funded Project for Interdisciplinary Exploration (No: TZJH2024027), the Natural Science Basic Research Program of Shaanxi (No: 2025JC-JCQN-094) and the Fundamental Research Funds for the Central Universities (No: QTZX25071).

### Conflict of Interest

none declared.

## Notes

https://doi.org/10.5281/zenodo.15637837

### Competing Interest Statement

The authors have declared no competing interest.

