## Supplementary information for "linearPOA: A parallel, memory-efficient framework for Partial Order Alignment with linear space complexity"


#### Illustration of quadratic space method of affine gap penalty POA

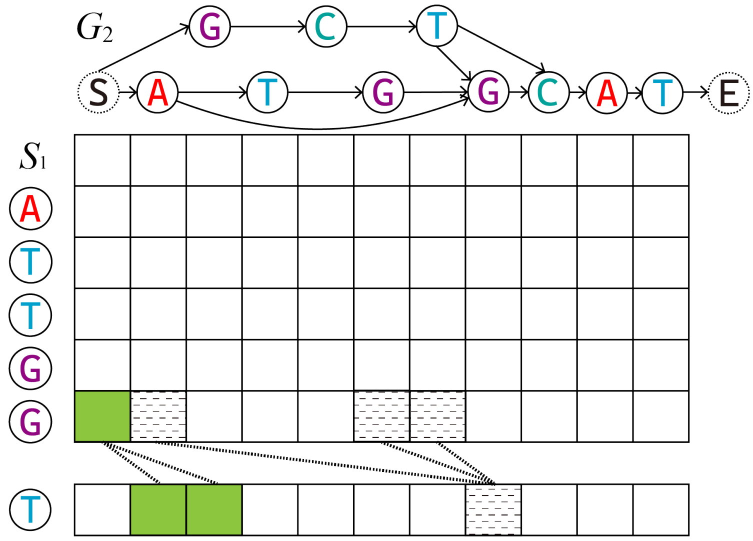


Figure S1. Illustration of quadratic space method of affine gap penalty POA. The matrix represents the alignment between sequence $S_1$ and graph $G_2$, where each cell corresponds to a state in the DP matrix. The dashed lines illustrate the recursive calculation flow that links subproblems based on the DP recurrence. A dashed line is included only when there is a valid edge between nodes in the DAG, reflecting the dependency between the preceding subproblem and the current one. For blurred cells in the last row, the algorithm considers the predecessor nodes in the previous row. These predecessor relationships are highlighted using dashed arrows. For colored cells, no predecessor nodes exist for the corresponding node $G_{2,j}$ in the graph. These cells are initialized using only the values in column 0, as indicated by the colored region in the matrix

### Pseudo code for linearPOA algorithm

Algorithm S1: Linear space complexity algorithm for affine gap penalty POA.

Input: Sequence $S_1$ and graph $G_2$, match $m$, mismatch $x$, gap open $o$, gap extension $e$
Output: The alignment result $R$ for $S_1$ and $G_2$
gap($n$) { if ($n$ == 0) return 0; else return $o + (n - 1)e$; }
chr_score($a, b$) { if ($a$ == $b$) return $m$; else return $x$;}
linearPOA_main($S_1, G_2, M, N$) {
  $R$ = [];
  linearPOA($S_1, G_2, S_{1,1}, S_{1,M}, M, G_{2S}, G_{2E}, N, o, o$);
  return sorted($R$);
}
linearPOA($S_1, G_2, s_a, s_b, l_1, g_x, g_y, l_2, tb, te$) {
  if ($l_1$ == 0) { // if $G_2$ has node, align the shortest path with gaps
    if ($l_2$ > 0) $R$.append(align(shortest_path($g_x, g_y$), gap));
  } else if ($l_2$ == 0) { // if no nodes in $G_2$, directly add sequences from $s_a$ to $s_b$
    if ($l_1$ > 0) $R$.append(align($s_a$ to $s_b$, gap));
  } else if ($l_1$ == 1) { // align sequence $s_a$ to graph nodes with $G_2$
    $g_{opt}$ = None; min_score = min($tb, te$) + $e$ + shortest_path_len($g_x, g_y$);
    for ($g_i$ in $G_2$) { // calculate the minimum score in $G_2$
      s = gap(shortest_path_len($g_x, g_i$) - 1) + chr_score($s_a, g_i$) + gap(shortest_path_len ($g_i, g_y$) - 1);
      // found better point, record the point and score
      if (s < min_score) { min_score = now_score; $g_{opt}$ = $g_i$; }
    }
    if ($g_{opt}$ != None) {
      $R$.append(align(shortest_path($g_x, g_{opt}$), gap));
      $R$.append(align($s_a, g_{opt}$));
      $R$.append(align(shortest_path($g_{opt}, g_y$), gap));
    } else {
      $R$.append(align(shortest_path($g_x, g_y$), gap));
      $R$.append(align($s_a$, gap));
    }
  } else { // divide the problem into subproblems
    $i^* = \lfloor l_1/2 \rfloor$;
    // return the $i^*$ line of $C$ and $D$ matrices, but the gap open value in row 0 of matrix $C$
    // is $tb/te$ (same as Myers' $O(N)$ space cost version (Myers and Miller, 1988))
    $\boldsymbol{CC}, \boldsymbol{DD}$ = DP($S_1, s_a, i^*, G_2, tb$); $\boldsymbol{CC}^r, \boldsymbol{DD}^r$ = DP($S_1', s_b, l_1 - i^*, G_2', te$);
    $j^* = \underset{0<j\leq l_2}{\operatorname{argmin}}\left(\boldsymbol{CC}[j] + \boldsymbol{CC}^r\left[N - next(g_j)\right], \boldsymbol{DD}[j] + \boldsymbol{DD}^r\left[N - next(g_j)\right] - g\right)$;
    if (($i^*, j^*$) is type 1) { // $j^*$ from $\boldsymbol{CC}[j] + \boldsymbol{CC}^r\left[N - next(g_{2,j})\right]$
      linearPOA($S_1, G_2, s_a, s_{i^*}, i^*, g_x, g_{j^*}, j^*, tb, o$);
      linearPOA($S_1', G_2', s_b, s_{i^*+1}, l_1 - i^*, g_y, next(g_{j^*}), l_2 - j^*, o, te$);

} else { // $j^*$ from $\boldsymbol{DD}[j] + \boldsymbol{DD}^r[N - next(g_j)] - g$

linearPOA($S_1, G_2, s_a, s_{i^*-1}, i^* - 1, g_x, g_{j^*}, j^*, tb, 0$);

$R$.append(align(($s_{i^*}, s_{i^*+1}$), gap));

linearPOA($S_1', G_2', s_b, s_{i^*+2}, l_1 - i^* - 1, g_y, next(g_{j^*}), l_2 - j^*, 0, te$);

}

}

}

#### Experimental Results on simulated dataset

To show the effect of different arguments between our methods and other methods, we tested abPOA, TSTA and our method. The result is shown in Table S1.

Table S1. Simulated dataset results. The argument of all test listed in this table are set as 0,6,5,3

| Test case | Technology | Depth | abPOA | | | TSTA | | | linearPOA | | |
|---|---|---|---|---|---|---|---|---|---|---|---|
| | | | FORAlign similarity | WFA2 similarity | minimap2 similarity | FORAlign similarity | WFA2 similarity | minimap2 similarity | FORAlign similarity | WFA2 similarity | minimap2 similarity |
| HIV | ont | 3 | 92.52% ±3.29% | 92.52% ±3.28% | 94.84% ±2.43% | 93.28% ±0.32% | 93.22% ±0.39% | 95.40% ±0.17% | 93.30% ±0.25% | 93.30% ±0.23% | 95.52% ±0.04% |
| | | 5 | 92.02% ±11.94% | 92.01% ±11.93% | 93.00% ±15.57% | 94.51% ±0.27% | 94.48% ±0.28% | 95.47% ±0.20% | 94.74% ±0.26% | 94.75% ±0.26% | 95.71% ±0.18% |
| | | 10 | 93.98% ±12.89% | 93.98% ±12.84% | 94.27% ±11.84% | 96.04% ±0.08% | 96.00% ±0.06% | 96.24% ±0.03% | 95.16% ±0.76% | 95.16% ±0.75% | 95.35% ±0.75% |
| | | 30 | 96.04% ±1.07% | 96.04% ±1.06% | 96.08% ±1.06% | 96.59% ±0.04% | 96.50% ±0.01% | 96.50% ±0.01% | 94.91% ±2.64% | 94.90% ±2.64% | 95.02% ±2.65% |
| | | 50 | 96.53% ±0.06% | 96.53% ±0.06% | 96.57% ±0.06% | 96.56% ±0.05% | 96.45% ±0.04% | 96.55% ±0.02% | 93.27% ±1.53% | 93.27% ±1.52% | 93.53% ±1.57% |
| | pacbio | 3 | 87.34% ±11.88% | 87.32% ±11.95% | 91.52% ±6.31% | 88.63% ±4.24% | 88.62% ±4.24% | 92.67% ±1.19% | 88.50% ±4.06% | 88.49% ±4.02% | 92.53% ±1.17% |
| | | 5 | 87.87% ±21.58% | 87.87% ±21.65% | 89.99% ±19.31% | 91.28% ±1.32% | 91.22% ±1.45% | 93.38% ±0.21% | 90.79% ±3.69% | 90.78% ±3.69% | 92.92% ±2.94% |
| | | 10 | 90.44% ±5.64% | 90.30% ±5.70% | 91.34% ±5.08% | 93.07% ±0.17% | 92.98% ±0.19% | 93.78% ±0.14% | 91.27% ±6.12% | 91.25% ±6.07% | 91.92% ±5.65% |
| | | 30 | 92.38% ±7.26% | 92.37% ±7.25% | 92.68% ±7.34% | 93.81% ±0.09% | 93.74% ±0.08% | 94.18% ±0.06% | 89.29% ±9.87% | 89.28% ±9.73% | 89.74% ±9.84% |
| | | 50 | 92.56% ±5.58% | 92.56% ±5.58% | 92.82% ±5.57% | 93.64% ±0.12% | 93.49% ±0.13% | 93.90% ±0.13% | 90.73% ±3.38% | 90.72% ±3.36% | 91.50% ±3.40% |
| mt | ont | 3 | 93.30% ±1.71% | 93.30% ±1.73% | 95.35% ±0.52% | 93.38% ±1.70% | 93.38% ±1.74% | 95.54% ±0.30% | 93.39% ±1.58% | 93.38% ±1.64% | 95.52% ±0.31% |
| | | 5 | 94.37% ±2.52% | 94.36% ±2.52% | 95.39% ±1.32% | 94.51% ±1.52% | 94.48% ±1.64% | 95.56% ±0.59% | 94.00% ±1.91% | 94.00% ±1.91% | 95.04% ±1.28% |
| | | 10 | 96.07% ±0.11% | 96.07% ±0.11% | 96.28% ±0.09% | 96.01% ±0.07% | 95.91% ±0.08% | 96.15% ±0.09% | 95.64% ±0.53% | 95.63% ±0.54% | 95.86% ±0.52% |
| | | 30 | 96.48% ±0.01% | 96.48% ±0.01% | 96.51% ±0.01% | 96.49% ±0.04% | 96.40% ±0.04% | 96.50% ±0.04% | 95.39% ±1.63% | 95.38% ±1.61% | 95.50% ±1.60% |
| | | 50 | 96.53% ±0.89% | 96.53% ±0.91% | 96.57% ±0.07% | 96.52% ±0.03% | 96.46% ±0.04% | 96.51% ±0.04% | 94.11% ±2.10% | 94.10% ±2.09% | 94.36% ±2.05% |
| | pacbio | 3 | 88.76% ±1.76% | 88.77% ±1.76% | 92.65% ±1.04% | 88.92% ±1.46% | 88.79% ±1.53% | 92.68% ±0.52% | 88.75% ±1.25% | 88.76% ±1.25% | 92.75% ±0.52% |
| | | 5 | 91.17% ±0.51% | 91.17% ±0.51% | 93.14% ±0.40% | 91.33% ±0.96% | 91.29% ±0.92% | 93.29% ±0.36% | 91.10% ±0.58% | 91.09% ±0.59% | 93.02% ±0.55% |
| | | 10 | 93.00% ±0.21% | 92.95% ±0.18% | 93.70% ±0.18% | 92.98% ±0.08% | 92.96% ±0.08% | 93.82% ±0.01% | 91.70% ±3.49% | 91.69% ±3.50% | 92.44% ±3.43% |
| | | 30 | 93.50% ±0.03% | 93.50% ±0.03% | 93.75% ±0.03% | 93.59% ±0.06% | 93.49% ±0.05% | 93.94% ±0.01% | 90.05% ±3.44% | 90.05% ±3.43% | 90.55% ±3.75% |
| | | 50 | 91.86% ±7.90% | 91.86% ±7.90% | 92.15% ±7.80% | 93.59% ±0.07% | 93.51% ±0.08% | 94.05% ±0.05% | 87.93% ±5.21% | 87.94% ±5.22% | 88.68% ±5.40% |

To Illustrate the speedup parallelization performance of linearPOA, we tested linearPOA using different numbers of threads and measured the corresponding execution time and memory usage. The results are shown in Figure S2 and Figure S3, covering all simulated datasets.

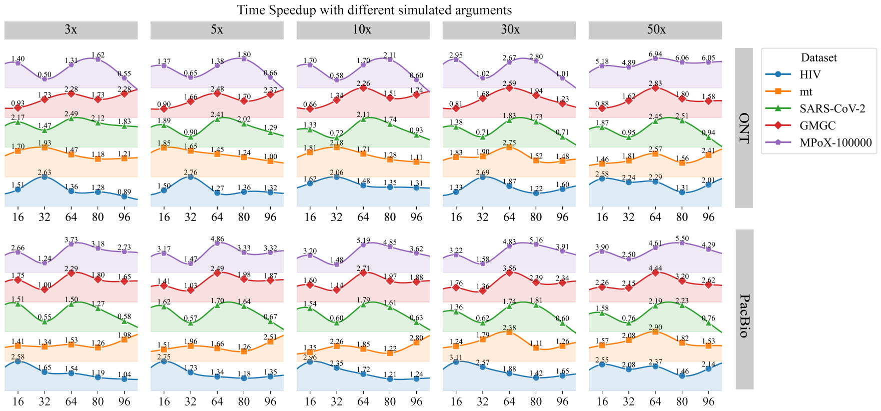


Figure S2. Experimental results between different threads in time speedup.

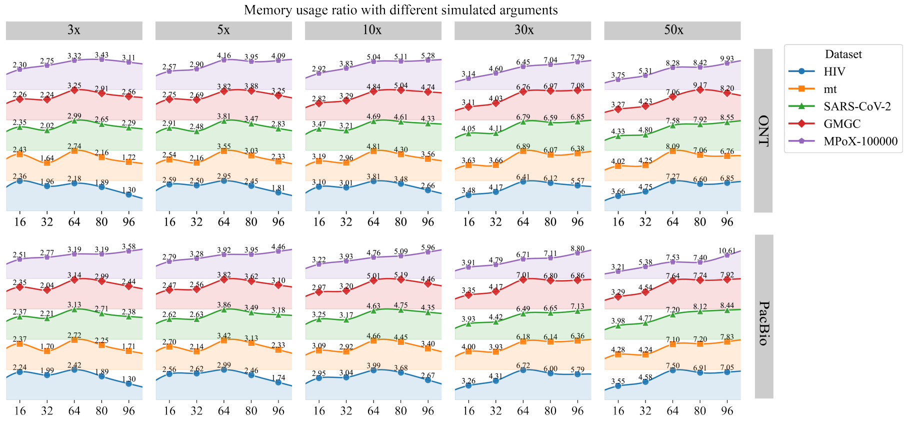


Figure S3. Experimental results between different threads in memory usage.

### Experimental Results on real dataset

Table S2 shows the detailed results in arguments 0,4,6,2. These methods are tested by the modified Racon.

Table S2. Detailed result for real datasets. These results are generated by arguments 0,4,6,2

| Test case | Technology | Depth | Window Size | Mapped Sequences | SPOA | | | | abPOA | | | | abPOA-band | | | |
|---|---|---|---|---|---|---|---|---|---|---|---|---|---|---|---|---|
| | | | | | Time /s | Memory /MB | Corrected Sequences similarity | Assebmled genome similarity | Time /s | Memory /MB | Corrected Sequences similarity | Assebmled genome similarity | Time /s | Memory /MB | Corrected Sequences similarity | Assebmled genome similarity |
| Lambda phage | ONT | 113 | 1000 | 515 | **28.88** | 452.71 | **88.67%±5.52%** | **97.56** | 166.47 | 488.24 | 87.55%±5.79% | 96.22 | 358.79 | 1361.42 | 83.22%±6.86% | 89.28 |
| | | | 2000 | 515 | **179.21** | 2361.52 | **90.41%±4.57%** | **97.67** | 440.8 | 2209.97 | 88.41%±5.26% | 96.34 | 941.27 | 5121.6 | 81.71%±7.17% | 80.74 |
| | | | 3000 | 515 | **240.5** | 4829.41 | **92.22%±3.33%** | **97.48** | 1100.54 | 6737.34 | 89.86%±4.16% | 91.86 | 1986 | 10178.78 | 81.41%±6.76% | 86.49 |
| | | | Window Size | Mapped Sequences | TSTA | | | | linearPOA | | | | | | | |
| | | | 1000 | 515 | 1122.2 | 415.06 | 75.52%±5.83% | 86.48 | 2104.42 | **176.35** | 72.64%±4.14% | 79.64 | | | | |
| | | | 2000 | 515 | 1949.22 | 1330.92 | 74.98%±6.53% | 87.73 | 4090 | **179.37** | 68.34%±5.46% | 86.24 | | | | |
| | | | 3000 | 515 | 2780.64 | 2766.75 | 72.34%±8.04% | 88.33 | 11914 | **182.25** | 67.97%±6.03% | 73.77 | | | | |
| Test case | Technology | Depth | Window Size | Mapped Sequences | SPOA | | | | abPOA | | | | abPOA-band | | | |
| E. coli K-12 | Hifi | 20 | 1000 | 5996 | **106.87** | 687.7 | **97.26%±1.16%** | 106.35 | 382.22 | 660.02 | 97.24%±1.12% | 98.71 | 198.01 | 671.61 | 96.90%±1.39% | 98.69 |
| | | | 2000 | 5998 | **204.6** | 974.73 | 95.18%±2.37% | 99.33 | 869.64 | 617.95 | 94.54%±2.87% | 95.39 | 719.4 | 1761.07 | 94.42%±2.98% | 97.4 |
| | | | 3000 | 6002 | **245.27** | 2193.88 | 92.36%±4.26% | 93.75 | 1151.44 | **605.89** | 91.84%±4.69% | 96.6 | 1460.51 | 5199.79 | 92.05%±4.55% | 96 |
| | | | Window Size | Mapped Sequences | TSTA | | | | linearPOA | | | | | | | |
| | | | 1000 | 5996 | 3674 | 927.75 | 95.74%±2.45% | **99.26** | 34410 | **657.66** | 84.74%±2.43% | 90.25 | | | | |
| | | | 2000 | 5998 | 6251 | 1720.5 | **97.14%±1.48%** | **99.39** | 29297 | **617.08** | 73.71%±5.79% | 82.82 | | | | |
| | | | 3000 | 6002 | 8690 | 3472.5 | **97.82%±1.06%** | **99.44** | 27323 | 625.88 | 65.90%±7.75% | 78.09 | | | | |
| Test case | Technology | Depth | Window Size | Mapped Sequences | SPOA | | | | abPOA | | | | abPOA-band | | | |
| E. coli K-12 | ONT | 54 | 1000 | 24165 | **1224.12** | 1844.8 | **96.67%±0.83%** | **98.3** | 6222 | 1403.29 | 96.74%±0.88% | 98.39 | 10466 | 2301.7 | 92.04%±3.05% | 96.97 |
| | | | 2000 | 24170 | 2289.88 | 3338.61 | **96.64%±0.93%** | 98.25 | 11184 | **1244.16** | 96.58%±1.04% | 98.26 | 24817 | 7332.44 | 87.49%±5.48% | 95.88 |
| | | | 3000 | 24172 | 4470 | 5646.58 | **96.64%±0.86%** | **98.42** | 16957 | 2720.92 | 96.27%±1.22% | 97.58 | 47187 | 16192.03 | 82.51%±7.55% | 94.23 |
| | | | Window Size | Mapped Sequences | TSTA | | | | linearPOA | | | | | | | |
| | | | 1000 | 24165 | 35806 | 1440.36 | 75.37%±3.62% | 84.95 | 327769 | 1417.41 | 76.75%±1.92% | 92.18 | | | | |
| | | | 2000 | 24170 | 65152 | 1950 | 76.70%±3.58% | 86.65 | 360289 | 1266.08 | 69.67%±3.83% | 94.87 | | | | |
| | | | 3000 | 24172 | 89040 | 3933.52 | 78.24%±3.33% | 87.23 | 415791 | **1229** | 67.85%±4.54% | 94.26 | | | | |
